# Spatial and temporal translocation of PKCα in single endothelial cell in response to focal mechanical stimulus

**DOI:** 10.1101/191833

**Authors:** Masataka Aarai, Toshihiro Sera, Takumi Hasegawa, Susumu Kudo

## Abstract

We observed the kinetics of protein kinase Cα (PKCα) and the intracellular Ca^2+^ wave in endothelial cells (ECs) in response to microscopic mechanical stress to investigate the effect of mechanical stress on PKCα translocation. The results show that a focal mechanical stimulus induced biphasic and directional PKCα translocation; PKCα initially translocated toward distinct spots near or at the membrane and then accumulated at the stimulus point. The low initial translocation occurred simultaneously in parallel with the increase in Ca^2+^. Initial translocation was inhibited in spite of Ca^2+^ increase when the diacylglycerol (DAG) binding domain of PKCα was inhibited, suggesting that translocation requires intracellular Ca^2+^ increase and DAG. On the other hand, high secondary translocation was delayed, occurring after the Ca^2+^ wave; however, this secondary translocation occurred even when Ca^2+^ release from the endoplasmic reticulum was inhibited, while it did not occur when the mechanosensitive (MS) channel was inhibited. These results indicated that at least Ca^2+^ influx through MS channels is required. Our results support the implication of PKCα in the Ca^2+^ signaling pathway in response to mechanical stress in ECs.

**Summary statement:** In response to a focal mechanical stimulus, PKCα in an endothelial cell was initially translocated toward distinct spots near or at the membrane and then accumulated at the stimulus point.

## Introduction

PKCα, which belongs to the conventional PKC family, plays an important role in vascular disease regulation, such as heart failure, atherosclerosis, endothelial cell (EC) proliferation, and endothelial barrier function (Konopatskaya and Poole, 2010). The cell membrane receptors are activated by various hormones and growth factors, resulting in the hydrolyzation of the membrane phospholipid phosphatidylinositol 4,5-bisphosphate (PIP2) by phospholipase Cβ (PLCβ) into inositol 1,4,5-triphosphate (IP3) and diacylglycerol (DAG). IP3 is released into the cytosol and acts as the intracellular store for Ca^2+^ release from the endoplasmic reticulum (ER). Finally, PKCα is activated and recruited to the plasma membrane by both C1 and C2 domains, which bind DAG and Ca^2+^, respectively.

ECs line the interior surface of blood vessels; therefore their functions are also regulated by various mechanical stresses, such as mechanical stretch and shear stress and pressure from the blood stream. Accordingly, the membrane receptors are activated by these mechanical stresses, and G-protein-coupled receptors (GPCRs) play crucial roles in the mechanical stress-induced signaling pathways (Li and Xu, 2000; Li and Xu, 2007). Furthermore, the mechanical stresses directly activate tyrosine kinase-coupled receptors (RTKs) in smooth muscle cells (SMC), including platelet-derived growth factor receptor (PDGF-R) (Li et al., 2003; Tanabe et al., 2000), resulting in the activation of PKCδ (Li et al., 2003), which is one of the novel PKCs and is activated by DAG alone. PDGF is activated by mechanical stress (Hu et al., 1998; Ma et al., 1999; Tanabe et al., 2000), and it activates PKCε, which is one of the novel PKCs, involving PLCγ (Moriya et al., 1996). While the intracellular Ca^2+^ increases in response to mechanical stretch via Ca^2+^-permeable mechanosensitive (MS) channel activation in human umbilical endothelial cells (Naruse and Sokabe, 1993), the inhibition of the MS channel prevents this Ca^2+^ wave from propagation. Tsukamoto et al. reported that PLC was activated in parallel with the intracellular Ca^2+^ wave in response to a focal mechanical stimulus (Tsukamoto et al., 2010). These previous results suggested that PKCα might be activated by mechanical stress. Indeed, previously, western blot analysis revealed that PKCα in SMC was shown to be activated by membrane stretch (Li et al., 2003).

Many pharmacological studies have shown that the translocation of conventional PKC to the membrane is observed in synchrony with the Ca^2+^ wave (Lenz et al., 2002; Oancea and Meyer, 1998; Schaefer et al., 2001; Sinnecker and Schaefer, 2004). However, the kinetics of PKC in response to mechanical stress have not yet been directly observed in living cells. In this study, we observed the kinetics of PKCα and the intracellular Ca^2+^ wave in bovine vascular ECs in response to microscopic mechanical stress to investigate the effect of mechanical stress on the translocation of PKCα. Additionally, we examined the effect of Ca^2+^ on the translocation of PKCα.

## Results

We first observed bovine aortic endothelial cells (BAECs) transfected with PKCα–Dronpa Green (DG) using confocal microscopy (Fig. 1). First we characterized the expression and localization of PKCα–DG in BAECs by western blot analysis and compared the results with those for wild-type PKCα (Fig. 1A and B). PKCα and DG were detected at approximately 80 kDa and 28.8 kDa, respectively, and therefore PKCα–DG was detected at 108.8 kDa. PKCα–DG was distributed homogenously based on the direct observation of DG fluorescence (Fig. 1C). PKCα was distributed in the cytosol based on the observation of anti-PKCα immunofluorescence (Fig. 1D). The colocalization of PKCα–DG fluorescence and anti-PKCα immunofluorescence demonstrated that the fusion protein was localized in the same area as PKCα (Fig. 1E). We also examined PKCα–DG translocation after the addition of ATP and PMA, which is known to activate PKCα, using confocal microscopy (Fig. 1F-I). Before the addition of ATP and PMA, PKCα–DG was distributed homogeneously throughout the cytoplasm, and absent in the nucleus. Stimulation with ATP and PMA caused the transient translocation of PKCα–DG to all parts of the plasma membrane and its depletion in the cytoplasm.

**Fig. 1.**
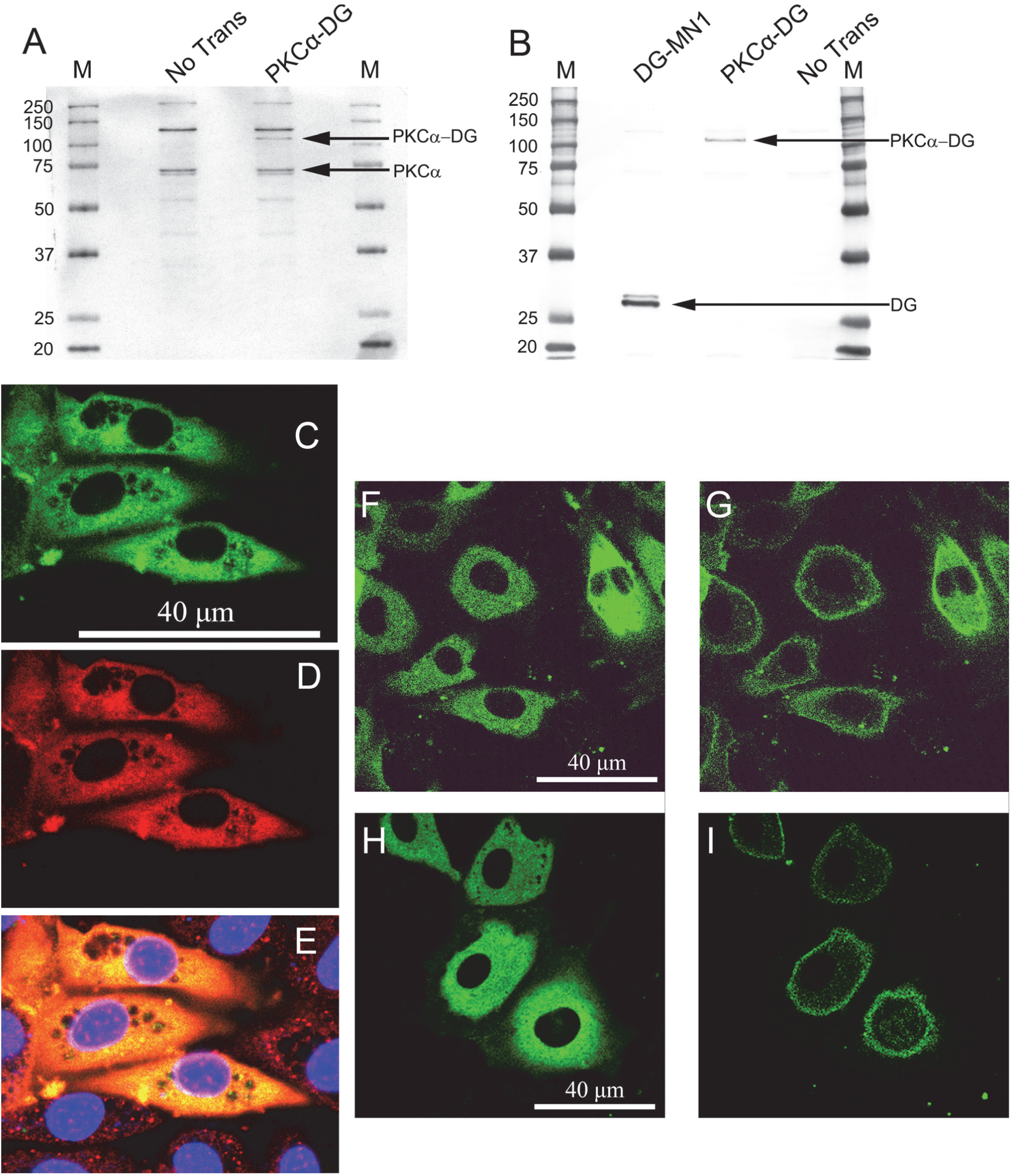
Characterization of the PKCα–DG expressed in bovine aortic endothelial cells. (A) Immunoblots of cell lysates from untransfected and PKCα–DG-BAECs probed with an antibody against PKCα. (B) Immunoblots of cell lysates from untransfected and PKCα–DG-BAECs probed with an antibody against DG. (C) PKCα-GFP. (D) Anti-PKCα. (E) Colocalization. (F and G)PKCα–GFP localization before (F) and 15 s after (G) stimulation with ATP. (H and I) PKCα–GFP localization before (H) and 15 min after (I) stimulation with PMA.

Figure 2 shows the representative intracellular Ca^2+^ wave (A) and PKCα translocation (B) in response to a focal mechanical stimulus. The fluorescent intensity of Ca^2+^ decreased at 10 s, indicating an increase in intracellular Ca^2+^ (Fig. 2A), and gradually returned to baseline (Fig. 2A). Regarding PKCα, the intensity increased gradually until 20 s, and then decreased. Although many pharmacological studies have revealed that PKCα was translocated to the plasma membrane due to a pharmacological stimulus, such as ATP (Fig. 1), our results suggested that PKCα was translocated to the stimulus point by a microscopic mechanical stimulus. Moreover, to analyze the translocation in detail, the image before stimulation (at 0 s in Fig. 2B) was subtracted from each image (Fig. 2B). The resulting image revealed that the fluorescent intensity was distributed at the membrane of the cell and, particularly, that it was not homogenous but rather at a distinct spot near or at the membrane close to the stimulus point (Fig. 2C). Fig. 2D shows the averaged fluorescent intensity ratio as a function of time at a distinct spot near the membrane, cytosol, and stimulus point. Especially, to demonstrate the inhomogeneous translocation of PKCα, the intensity ratios at the membrane were presented at points both near and far from the stimulus point. Ca^2+^ intensity decreased gradually and reached the minimum at approximately 7.5 s after stimulus, and then gradually increased. Compared among regions, Ca^2+^ intensity changed at the stimulus point, in the cytoplasm, and then at the membrane. PKCα intensity decreased soon after the stimulus since the membrane was pushed by the mechanical stimulus, which was observed in the previous study (Tsukamoto et al., 2010). Thereafter, it increased gradually at the membrane, particularly near the stimulus point, and at the stimulus point, but not in the cytosol. At the membrane near the stimulus point, the intensity reached its maximum at 9.2 ± 0.6 s (mean ± SE), and decreased gradually and returned to base level at approximate 30 s. On the other hand, the intensity continued to increase at the stimulus point after it had reached the maximum of the surrounding membrane, reaching a maximum at 21.7 ± 1.5 s (mean ± SE). Finally, it returned to base level at approximate 80 s. The peak intensities at the stimulus point and at the membrane near the stimulus point were 2.60 ± 0.10 and 1.62 ± 0.05 (mean ± SE), respectively. The peak time and intensity were significantly delayed and were higher at the stimulus point than at the membrane. Here we refer to PKCα translocation at a distinct spot near to the membrane and at the stimulus point as the initial and secondary translocations, respectively. The initial translocation occurred simultaneously with Ca^2+^ increase; however, the secondary translocation occurred after the Ca^2+^ activity, which is not consistent with previous pharmacological reports (Lenz et al., 2002; Oancea and Meyer, 1998; Schaefer et al., 2001; Sinnecker and Schaefer, 2004).

**Fig. 2.**
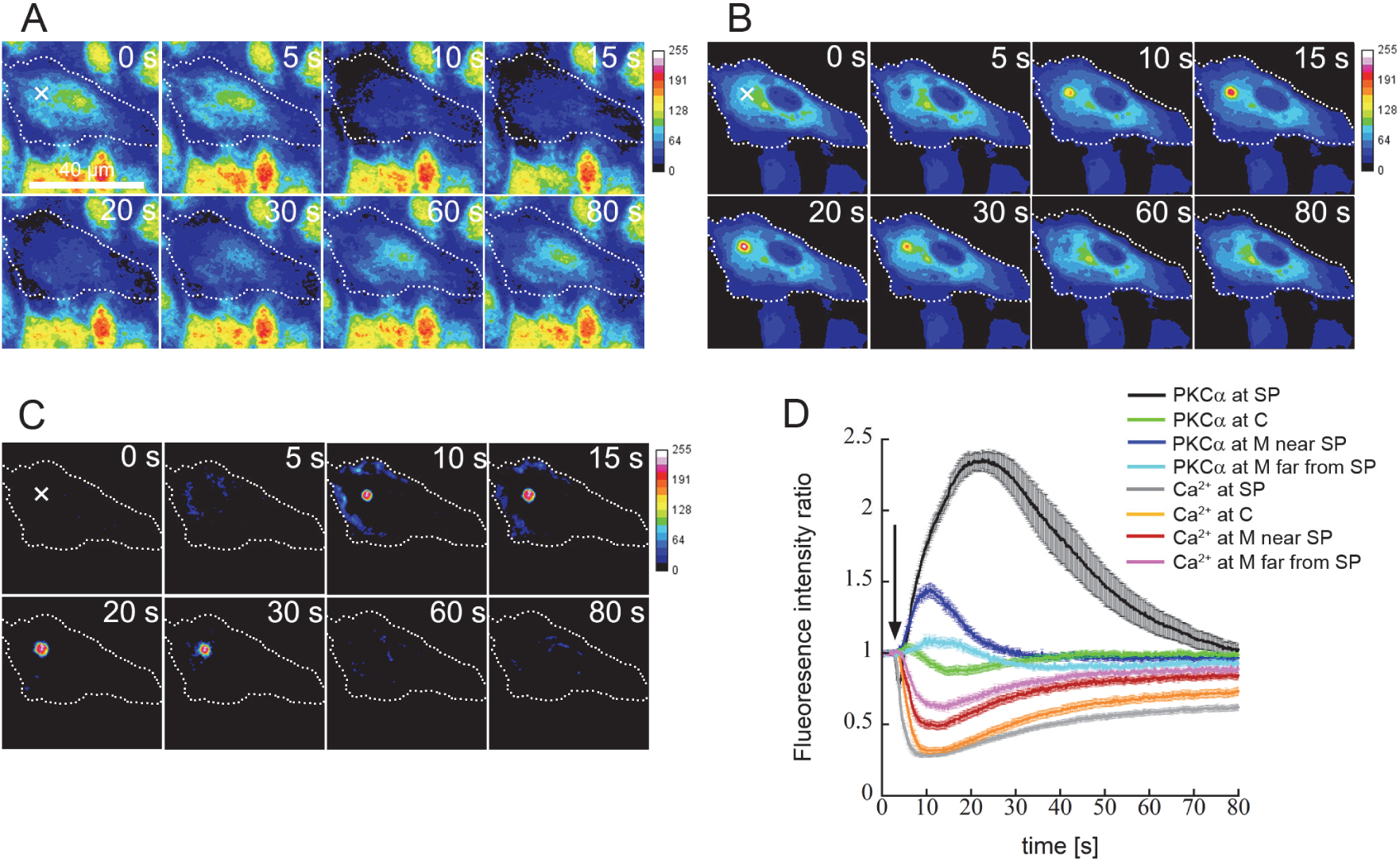
The Ca^2+^ wave and PKCα kinetics under the mechanical stimulus. The representative intracellular Ca^2+^ wave (A) and PKCα translocation (B and C). (C) is normalized PKCα–DG translocation, which is obtained by subtracting the images at *t* = 0 from every time-series image in (B). The cell geometry and stimulus point are indicated by the dotted line and “×”, respectively. The intensities of PKCα–DG and Ca^2+^ are shown using a pseudo color scale. (D) is the fluorescent intensity ratio of calcium and PKCα-DG at the stimulus point (SP), membrane (M), and cytoplasm(C) (averaged values ± S.E., N = 9, n = 17). PKCα translocations at M were presented near and far from SP, respectively. The cells are stimulated at 3 s (arrow).

Figure 3 shows the representative intracellular Ca^2+^ wave (A) and PKCα translocation (B) under the inhibition of C1 domains of PKCα. PKCα was not activated and translocated toward the membrane and at stimulus point even though a Ca^2+^ wave was observed (Fig. 3C). Figure 4 shows the representative intracellular Ca^2+^ wave (A) and PKCα translocation (B) under the inhibition of C2 domains of PKCα. Figure 4C shows the resulting image of PKCα after subtracting the image before stimulation, and PKCα translocated toward the membrane. Figure 4D shows the averaged fluorescent intensity ratio as a function of time. In contrast to the inhibition of its translocation under the C1 domain, PKCα was translocated to a localized spot near membrane; however, the fluorescent intensity did not return to base line at 80 s. PKCα intensity at the stimulus point did not increase, and the secondary translocation at the stimulus point was not observed.

**Fig. 3.**
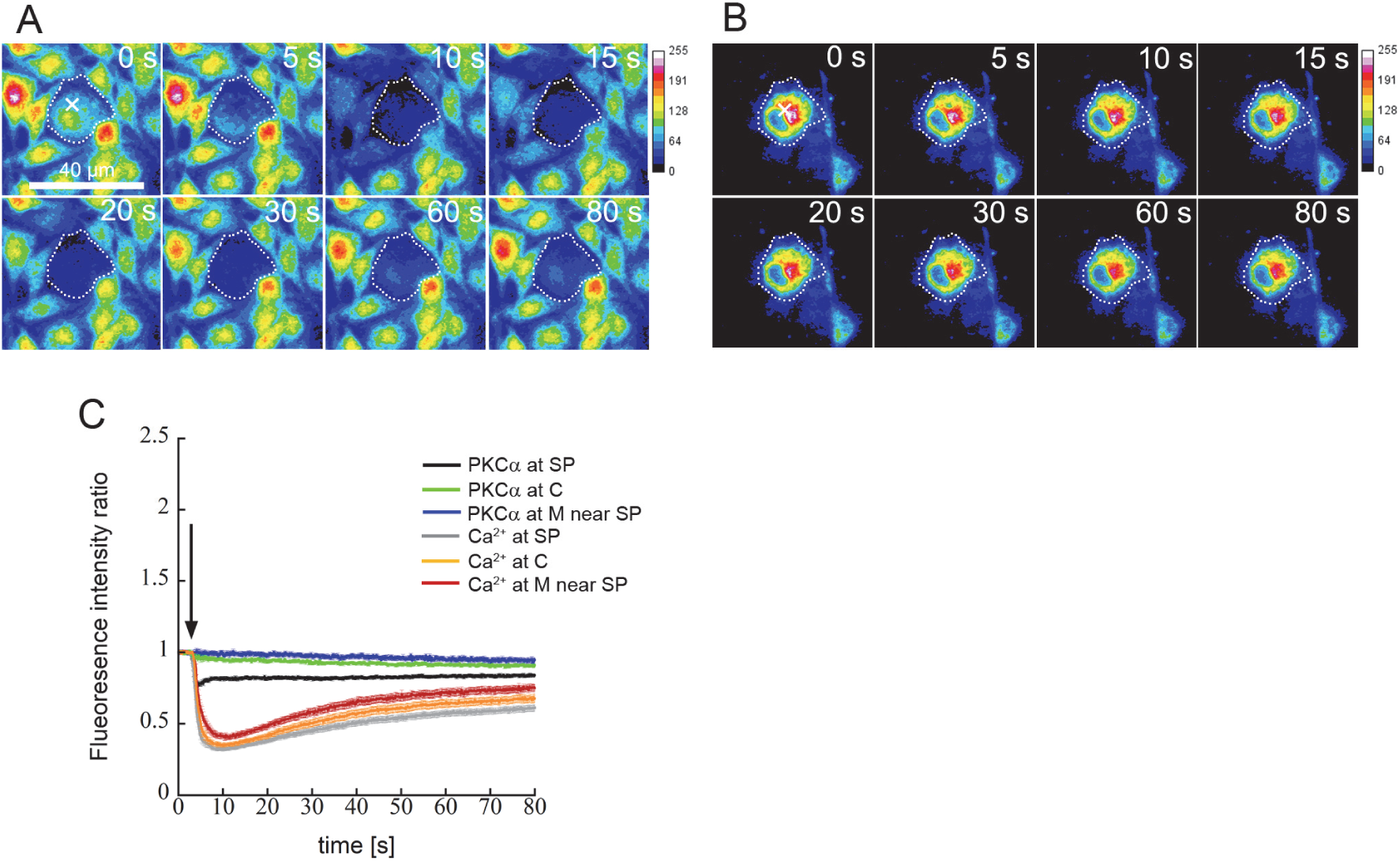
The Ca^2+^ wave and PKCα kinetics under the mechanical stimulus when the C1 domain of PKCα was inhibited. The representative intracellular Ca^2+^ wave (A) and PKCα translocation (B). The cell geometry and stimulus point are indicated by the dotted line and “×”, respectively. The intensities of PKCα–DG and Ca^2+^ are shown using a pseudo color scale. (C) shows the fluorescent intensity ratio of calcium and PKCα-DG at the stimulus point (SP), membrane (M), and cytosol (C) (averaged values ± S.E., N = 6, n = 20). The cells are stimulated at 3 s (arrow).

**Fig. 4.**
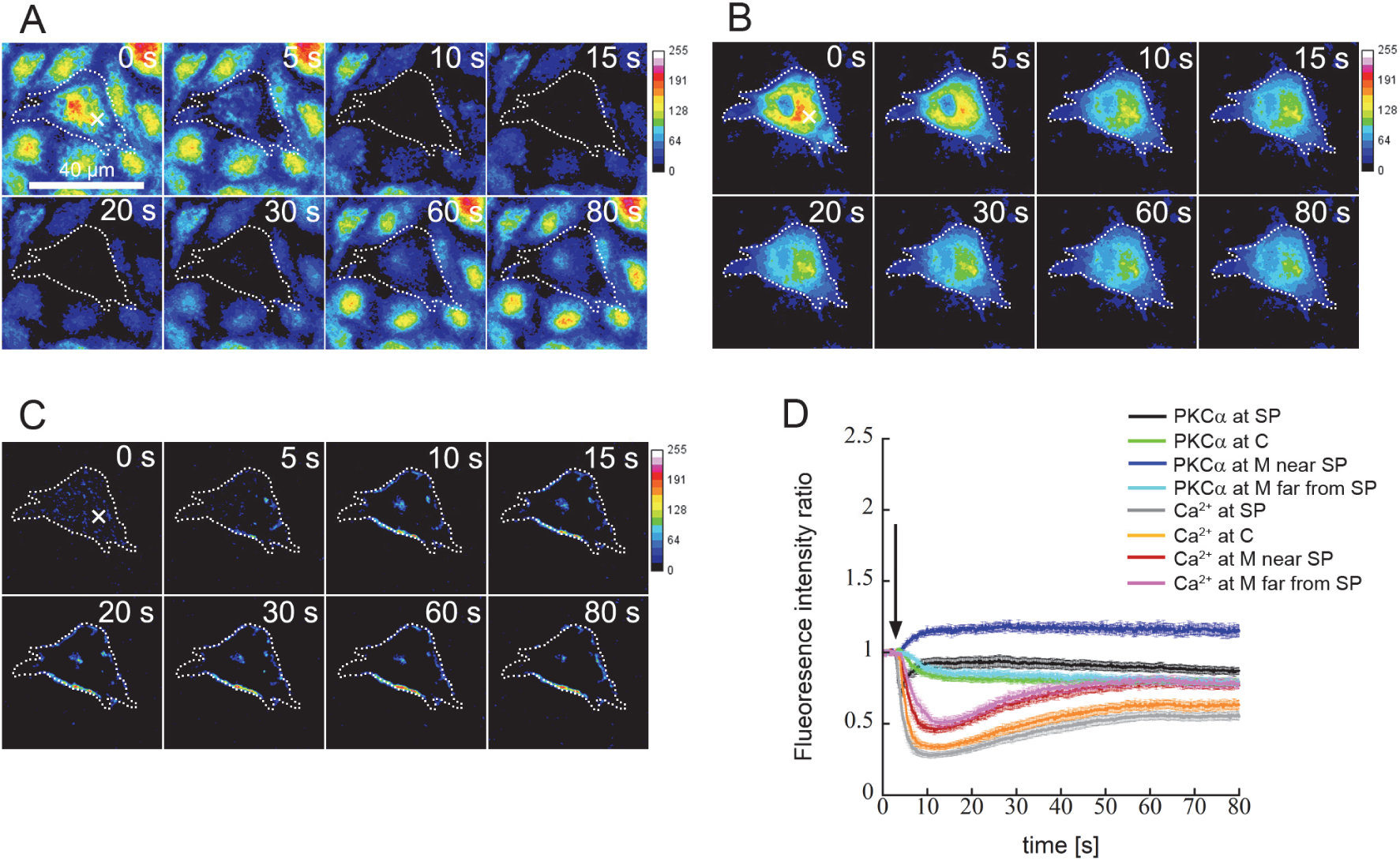
The Ca^2+^ wave and PKCα kinetics under the mechanical stimulus when the C2 domain of PKCα was inhibited. The representative intracellular Ca^2+^ wave (A) and PKCα translocation (B and C). (C) is normalized PKCα–DG translocation, which is obtained by subtracting the images at *t* = 0 from every time-series image in (B). The cell geometry and stimulus point are indicated by the dotted line and “×”, respectively. The intensities of PKCα–DG and Ca^2+^ are shown using a pseudo color scale. (D) is the fluorescent intensity ratio of calcium and PKCα-DG at the stimulus point (SP), membrane (M), and cytoplasm (C) (averaged values ± S.E., N = 5, n = 15). PKCα translocations at M were presented near and far from SP, respectively. The cells are stimulated at 3 s (arrow).

Figure 5 shows the representative intracellular C^2+^ wave (A) and PKCα translocation (B) under the inhibition of Ca^2+^ release from the ER. Ca^2+^ intensity changed in the stimulated cell only, and not in the neighboring cells. Figure 5C shows the subtracted image of PKCα using the image before stimulation, and PKCα translocated toward the membrane, particularly near the stimulus point. Figure 5D shows the averaged fluorescent intensity ratio of Ca^2+^ and PKCα-DG. Because of the Ca^2+^influx from the extracellular medium, a Ca^2+^ wave was observed and the maximum intensity at the stimulus point did not differ significantly from that of the control (Fig. 2). Regarding PKCα kinetics, the intensity reached its maximum (2.11 ± 0.07, mean ± SE), (1.52 ± 0.07, mean ± SE) at 21.9 ± 2.0 s and 7.8 ± 0.7 s at the stimulus point and at the membrane near the stimulus point, respectively. The peak time and intensity were significantly delayed and higher at the stimulus point than at the membrane, suggesting that both initial and secondary translocations were exhibited as control group (Fig. 2). Moreover, the initial translocation also occurred simultaneously with the Ca^2+^ increase. On the other hand, when the Ca^2+^ influx from the extracellular space was inhibited by the MS channel inhibitor (Fig. 6) and the removal of extracellular Ca^2+^ (Fig. 7), PKCα did not translocate at the stimulus point. particularly, both the Ca^2+^ wave and the PKCα translocation were not observed under the MS channel inhibitor (Fig. 6). The initial decrement due to the mechanical push was observed only when the MS channels were inhibited (Fig. 6). When the extracellular Ca^2+^ was removed (Fig. 7), Ca^2+^ increased because Ca^2+^ was released from the ER. However, the maximum value was not significantly lower than that at the control (Fig. 2) and under the inhibition of Ca^2+^ release from the ER (Fig. 5). Figure 7C shows the image of PKCα after subtracting the image before stimulation. PKCα intensity at the membrane near the stimulus point increased and reached its maximum (1.38 ± 0.05, mean ± SE) only at 8.1 ± 1.3 s, indicating that the initial translocation of PKCα only was exhibited when the extracellular Ca^2+^ was removed. This initial translocation also occurred simultaneously with Ca^2+^ increase.

**Fig. 5.**
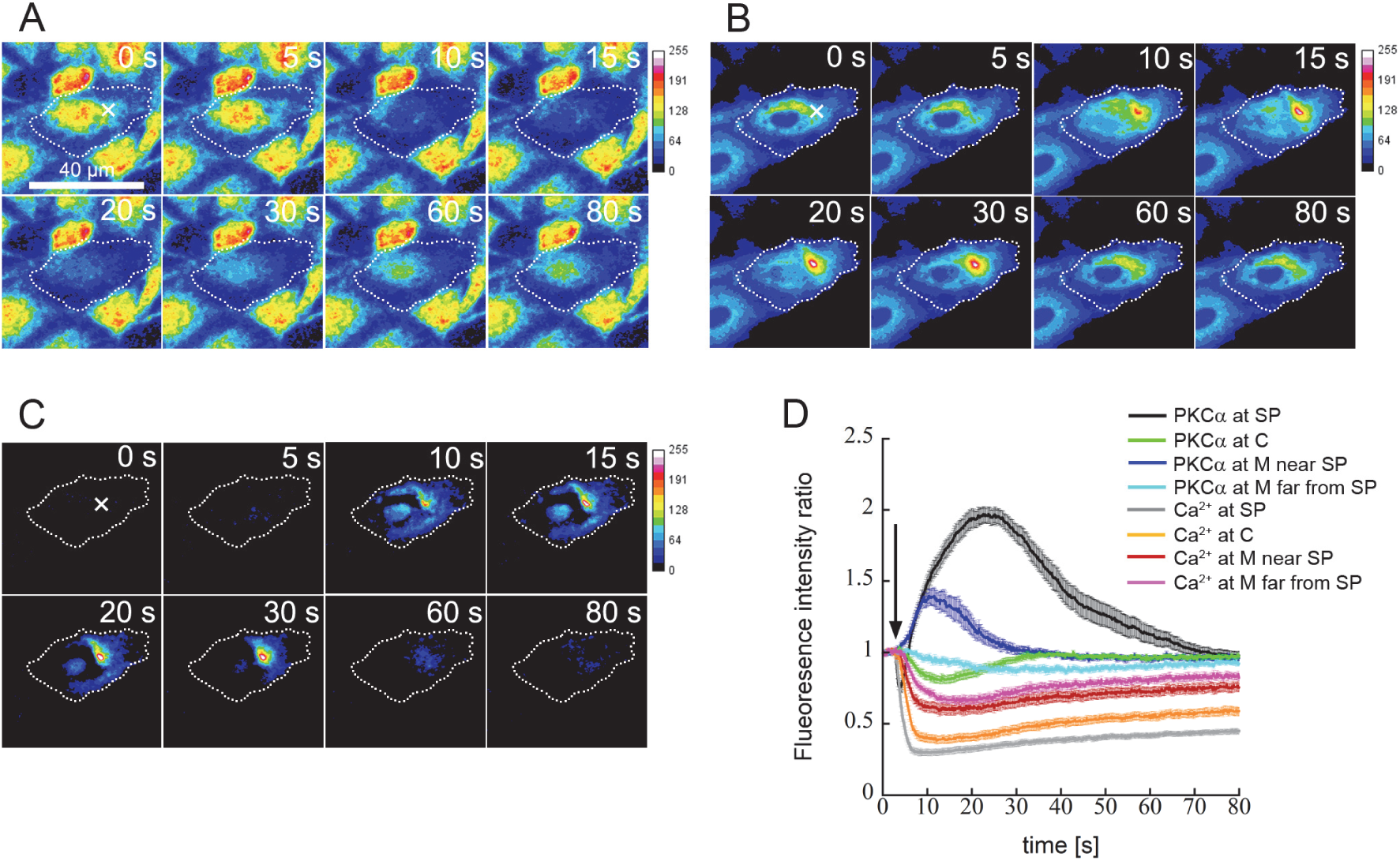
The Ca^2+^ wave and PKCα kinetics under the mechanical stimulus when Ca^2+^ release from the endoplasmic reticulum (ER) was inhibited. The representative intracellular Ca^2+^ wave (A) and PKCα translocation (B and C). (C) is normalized PKCα–DG translocation, which is obtained by subtracting the images at *t*= 0 from every time-series image in (B). The cell geometry and stimulus point are indicated by the dotted line and “×”, respectively. The intensities of PKCα–DG and Ca^2+^ are shown using a pseudo color scale. (D) shows the fluorescent intensity ratio of calcium and PKCα-DG at the stimulus point (SP), membrane (M), and cytosol (C) (averaged values ± S.E., N = 5, n = 13). PKCα translocation at M was presented near and far from SP, respectively. The cells are stimulated at 3 s (arrow).

**Fig. 6.**
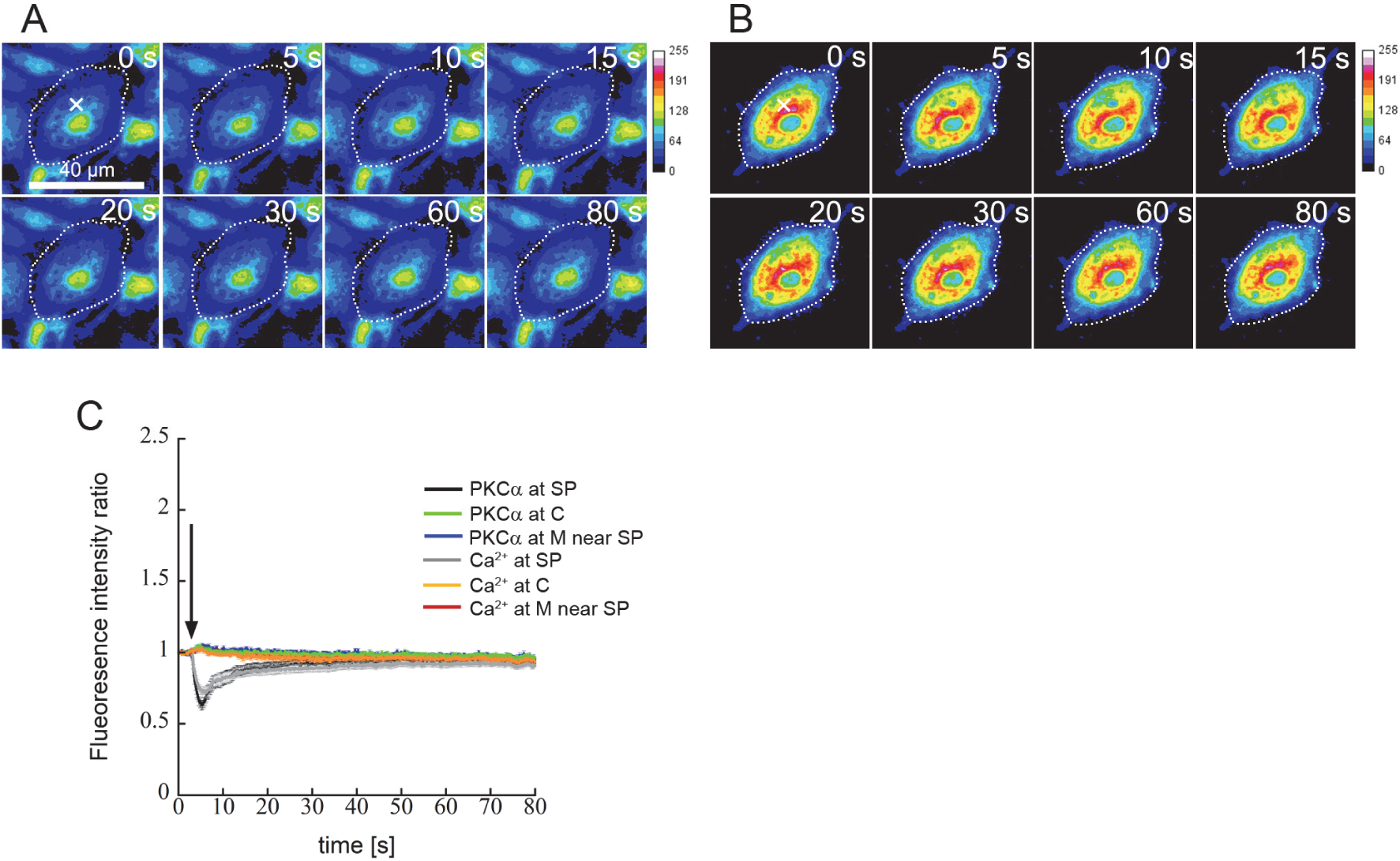
The Ca^2+^ wave and PKCα kinetics under the mechanical stimulus when the MS channel was inhibited. The representative intracellular Ca^2+^ wave (A) and PKCα translocation (B). The cell geometry and stimulus point are indicated by the dotted line and “×”, respectively. The intensities of PKCα–DG and Ca^2+^ are shown using a pseudo color scale. (C) shows the fluorescent intensity ratio of calcium and PKCα-DG at the stimulus point (SP), membrane (M), and cytoplasm (C) (averaged values ± S.E., N = 5, n = 16). The cells are stimulated at 3 s (arrow).

**Fig. 7.**
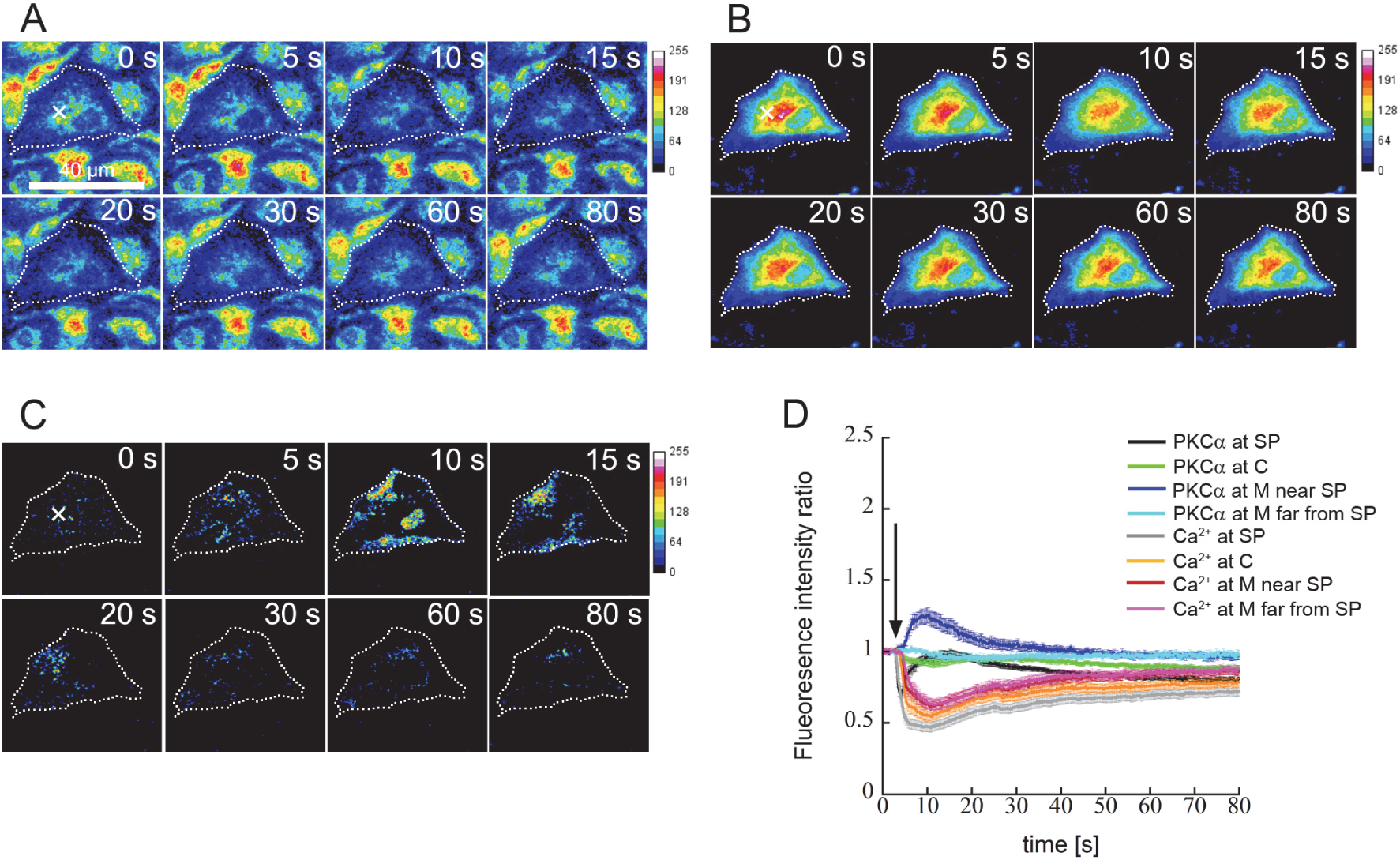
The Ca^2+^ wave and PKCα kinetics under the mechanical stimulus when the extracellular Ca^2+^ was removed. The representative intracellular Ca^2+^ wave (A) and PKCα translocation (B and C).(C) is the normalized PKCα–DG translocation, which is obtained by subtracting the images at *t* = 0 from every time-series image in (B). The cell geometry and stimulus point are indicated by the dotted line and “×”, respectively. The intensities of PKCα–DG and Ca^2+^ are shown using a pseudo color scale. (D) shows the fluorescent intensity ratio of calcium and PKCα-DG at the stimulus point (SP), membrane (M), and cytoplasm (C) (averaged values ± S.E., N = 5, n = 16). PKCα translocations at M were presented near and far from SP, respectively. The cells are stimulated at 3 s (arrow).

## Discussion

PKCα is activated by both Ca^2+^ and DAG, and the hallmark of its activation is its translocation from the cytosol to the plasma membrane (Newton, 2001). In particular, PKCα is translocated homogenously to the membrane under pharmacological agents, as shown in Fig. 1. On the other hand, previous studies using physiological agonists reported PKCα translocation is regulated by temporal and spatial changes in intracellular Ca^2+^ and that PKCα is translocated to different areas of the cell membrane depending on the Ca^2+^ influx from extracellular space (Maasch et al., 2000). In this study, we investigated PKCα translocation in the ECs in response to a microscopic mechanical stimulus using a DG fusion protein which exhibited the properties of the intact PKCα. Our results show that a focal mechanical stimulus induced a biphasic and directional PKCα translocation; namely, PKCα initially translocated toward distinct spots near or at the membrane, especially near the stimulus point, and then accumulated at the stimulus point. The low initial translocation occurred simultaneously with the increase in intracellular Ca^2+^, while the high secondary translocation was delayed, occurring after the Ca^2+^ wave, it required Ca^2+^ influx through MS channels.

A focal mechanical stimulus induced the microscopic mechanical stress on the cell membrane. Several membrane proteins, such as growth factor receptors, GPCRs, ion channels, caveolins and pumps could be candidate mechanosensors in the membrane (Li and Xu, 2007). Mechanical stress activates RTKs including PDGF-R, which regulates PKCδ, leading to the migration of smooth muscle cell (Li et al., 2003). On the other hand, G proteins, which include small molecular G proteins and heterotrimeric G proteins, could be activated by mechanical stress (Erickson et al., 2001; Gudi et al., 1998; Ishida et al., 1997; Yellowley et al., 1999) and implicated in the mechanical stress-induced signal pathways (Li and Xu, 2007). These candidate mechanosensors may be essential in biphasic PKCα translocation in response to mechanical stress. Maasch et al. demonstrated that localized Ca^2+^ determined the spatial and temporal PKCα translocation in rat vascular smooth muscle cell, and reported that thrombin acting on GPCRs induced rapid PKCα translocation; on the other hand, PDGF acting on PGDF-R in RTKs induced slow PKCα translocation (Maasch et al., 2000). Thrombin induced a PKCα translocation to spots near the membrane at 30 s and was reversible after 2 min, while PDGF induced a translocation to the membrane within 60 s and was reversible after 8–9 min (Maasch et al., 2000). Additionally, thrombin- and PDGF-induced translocation of PKCα require different Ca^2+^resources; thrombin-induced translocation is dependent on Ca^2+^ release from the ER and PDGF-induced translocation is dependent on Ca^2+^influx from the extracellular space (Maasch et al., 2000). Indeed, our results show that the initial translocation—but not the secondary translocation—was observed when extracellular Ca^2+^was removed (Fig. 7), and that secondary translocation was observed in spite of the inhibition of Ca^2+^ release from the ER (Fig. 5). Even when Ca^2+^ release from the ER was inhibited, initial translocation was observed (Fig. 5), and possibly Ca^2+^ influx from extracellular space may be substitute for Ca^2+^ release from ER. Our results suggested that both the initial and the secondary translocations require Ca^2+^ increase, and for the secondary translocation specifically, Ca^2+^ influx from the extracellular space is necessary. Since both GPCRs and RTKs are activated by mechanical stretch stresses (Hu et al., 1998; Li and Xu, 2007; Li et al., 2003), the focal mechanical stimulus in this study may activate both receptors microscopically. GPCRs and RTKs may be implicated in both the initial and the secondary translocations independently, resulting in the different activation timing of PKCα.

Previously, the translocation kinetics of PKC were parameterized by fitting a modified Bateman equation, as follows (Schaefer et al., 2001; Sinnecker and Schaefer, 2004):

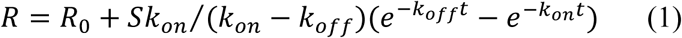

where *R* is the ratio of fluorescenct intensity at the translocation region (accumulated distinct point near the membrane or stimulus point) to the non-translocation region (cytoplasm), *R0* is the ratio in resting cell just before stimulus, *S* is the constant, and *t* is time. The terms *kon* and *koff* are the rate constants for association and dissociation, respectively. In this study, the initial translocation was observed under the control and inhibition of Ca^2+^ release from the ER, and the removal of extracellular Ca^2+^, and the secondary translocation was observed under the control and inhibition of Ca^2+^ release from the ER; therefore, these phenomena were quantified by fitting each translocation kinetic to Eqn 1 (Table 1). Regarding the initial translocation, there were no significant differences between the three conditions, and PKCα was activated with an increase in Ca^2+^ concentrations. PKCα repetitively translocated during Ca^2+^ oscillation with a tight coupling to the frequency of Ca^2+^ waves in human embryonic kidney cells under 20 μM carbachol (Schaefer et al., 2001). Moreover, there were no significant differences in *kon* and *koff*, and the initial translocation exhibited *kon*≅*koff* under these conditions, with ranges of 0.17 s^-^1^^–0.20 s^-^1^^ and 0.16 s^-^1^^–0.18 s^-^1^^, respectively. These results are similar to those using pharmacological agents (Sinnecker and Schaefer, 2004), suggesting that the initial translocation in response to a focal mechanical stimulus may have the same mechanism as pharmacological agents involving GPCRs. Although extracellular Ca^2+^ was chelated by exchanging the extracellular medium for medium containing EGTA, the intracellular Ca^2+^ wave was nevertheless observed (Fig. 7B), which is consistent with the previous study of PLC activation in mechanical simulation (Tsukamoto et al., 2010). This PLC activation was involved in the Ca^2+^ release from the ER and the Ca^2+^ wave in response to the mechanical stimulus (Tsukamoto et al., 2010), suggesting that DAG may be also produced. Furthermore, despite the Ca^2+^ wave, the initial translocation was not observed under the inhibition of the DAG-dependent C1 domain of PKCα (Fig. 3). These results suggested that this initial translocation required an increase in Ca^2+^ and the DAG-dependent C1 domain of PKCα. In contrast, there was no significant difference in *kon* and *koff* of the secondary translocation between two conditions, and the values were significantly smaller than those of the initial translocation, indicating that mechanics of the each translocation might be different.

**Table 1.** The association (*k*_*on*_) and dissociation (*k*_*off*_) of initial and secondary translocations of PKCα. Data are averaged values ± S.E. ^#^*P* < 0.05 vs initial translocation.

|  | Initial translocation |  | Secondary translocation |  |
| --- | --- | --- | --- | --- |
| | $k_{on}$ | $k_{off}$ | $k_{on}$ | $k_{off}$ |
| Control | $0.180 \pm 0.012$ | $0.181 \pm 0.011$ | $0.084 \pm 0.006^{\#}$ | $0.084 \pm 0.006^{\#}$ |
| Inhibition of Ca<sup>2+</sup> release from ER | $0.168 \pm 0.010$ | $0.164 \pm 0.011$ | $0.082 \pm 0.008^{\#}$ | $0.087 \pm 0.010^{\#}$ |
| Removal of extracellular Ca<sup>2+</sup> | $0.197 \pm 0.015$ | $0.161 \pm 0.022$ | - | - |

In this study, in response to microscopic mechanical stress, initial translocation was toward distinct spots near or at the plasma membrane—particularly near the stimulus point—and secondary translocation was at the stimulus point. As discussed above, the initial and secondary translocations by physiological agonists may be related to GPCR and TRK activation, respectively, and particularly PKCα in the secondary translocation might be accumulated at the membrane by TRKs activation. Conversely, Maasch et al. reported the local Ca^2+^ changes affected PKCα kinetics and proposed that the local Ca^2+^ increased the affinity of binding sites for PKCα membrane for the possible mechanism (Maasch et al., 2000). A small increase in Ca^2+^ increases the interaction between PKCα and the membrane, and a further increase would lead to PKCα activation in these areas (Luo and Weinstein, 1993; Maasch et al., 2000). Medkova and Cho proposed that the conventional PKC initially binds to the membrane surface via the Ca^2+^-dependent C2 domain, and then undergoes conformational changes that include the insertion of a C1a domain, which is one component of the C1 domain, into the membrane, resulting in the optional DAG binding (Medkova and Cho, 1999). Nalefski and Newton also proposed that the binding of Ca^2+^ to the C2 domain leads to a significantly higher affinity for membranes and stabilizes the C2 domain–membrane complex, and then searches two-dimensional space for DAG as long as the Ca^2+^ remain high (Nalefski and Newton, 2001). Based on these mechanisms, since the initial translocation is triggered by the Ca^2+^ wave across a cell, PKCα could translocate toward the membrane. However, DAG production would be lower when triggered by a focal mechanical stress than by pharmacological agonists, resulting in its translocation not being homogenous at the membrane and at localized spots near the stimulus point at the membrane. PLC activation occurred in broader cytosolic regions in parallel with the Ca^2+^ wave in response to mechanical stimulus, and particularly the high activation was localized at the membrane near the stimulus point(Tsukamoto et al., 2010). Our results show that the intensity of PKCα-DG remained relatively constant or decreased at the membrane far from the stimulus point in spite of the Ca^2+^ increase (Fig. 2D, 5D, and 7D). On the other hand, since the secondary translocation is dominated by the Ca^2+^ influx from the extracellular space, PKCα could be accumulated at the focal mechanical spot.

PKCα was translocated toward the membrane in response to mechanical stress despite the inhibition of the C2 domain, and this activation endured for longer than the other initial translocation (Fig. 4). Previous studies report that mechanical stress activates PLC (Tsukamoto et al., 2010) and may produce DAG, and deletion of the C2 domain of PKCα abolishes the translocation by ionomycin (Maasch et al., 2000). Therefore, this mechanism should not be the same as the initial translocation, in which both Ca^2+^and DAG may be involved, but may instead be dominated only by DAG. PKCα is activated directly by phorbol ester, which is analogous to DAG, and this mechanism might be different from the initial translocation (Medkova and Cho, 1999). As the different mechanism, reversible and irreversible activations of PKC were proposed. Based on this mechanism, phorbol ester induced sustained activation of PKC by forming an irreversible PKC membrane complex (Kazanietz et al., 1992). On the other hand, DAG acted through a reversible complex (Bazzi and Nelsestuen, 1989) and its activation was short-lived because DAG was rapidly metabolized (Newton, 2001); both, however, caused a dramatic increase in the affinity of PKC for membranes, serving as “molecular glue” to recruit PKC to the membrane (Newton, 2001). The resulting, activation time in response to mechanical stress may be longer than calcium-related initial translocation.

Initial and secondary translocations exhibited *kon* ≅ *koff*. Sinnecker and Schaefer proposed that *kon* ≅ *koff* implies a lower DAG concentration, while *kon* > *koff* implies a higher DAG concentration, and DAG binding delays the dissociation (Sinnecker and Schaefer, 2004). Regarding the initial translocation, as discussed above, DAG might be produced through PLC activation (Tsukamoto et al., 2010). The secondary translocation was inhibited when the C1 domain was inhibited, suggesting that secondary translocation may require DAG. However, since DAG is rapidly metabolized (Newton, 2001), it remains unclear whether secondary translocation requires DAG and whether DAG is also accumulated at the mechanical stress spot. Once PKC binds to the membrane via the C2 domain, it inserts a C1a domain for searching for DAG and releases the pseudo-substrate region from the active site, resulting in PKCα activation (Medkova and Cho, 1999). Therefore, the C2 domain of PKCα alone can bring PKC molecules to the membrane, and the membrane binding of the C1 domain might be primed by the membrane binding of the C2 domain (Kazanietz et al., 1992; Medkova and Cho, 1999).

From our results, it also remains unclear whether GPCRs and RTKs are activated directly by a focal mechanical stress. Previous studies reported that ATP release into extracellular spaces is induced by a mechanical stretch (Grygorczyk et al., 2013; Takahara et al., 2014). Takada et al. reported that this extracellular ATP is released through mechanosensitive hemichannels and acts as an autocrine mediator to activate GPCRs (Takada et al., 2014). On the other hand, Hayakawa et al. demonstrated that the increase in tension of actin stress fibers could activate MS channels and the actin skeleton acted as a force-transmitting and–focusing device between integrin and MS channels using ECs (Hayakawa et al., 2008). Indeed, in this study, the Ca^2+^ wave and PKCα translocation were not observed when MS channels were inhibited (Fig. 6), suggesting that GPCRs and RTKs might be activated by a focal mechanical stress via MS channel activation.

In this study, a focal mechanical stimulus induced the biphasic and directional PKCα translocation in single ECs. In its initial translocation that may involve the GPCRs signal pathway, PKCα is translocated toward the membrane in parallel with the Ca^2+^ wave. In secondary translocation, which may involve the RTKs signal pathway, PKCα was accumulated to the focal stimulus point after the Ca^2+^wave. However, Ca^2+^influx through MS channels from extracellular space, not Ca^2+^ release from ER, is required. PKCα is involved in several physiological functions, in the migration, proliferation, and angiogenesis of ECs (Bokhari et al., 2006; Newton, 2001; Wang et al., 2002); therefore, PKCα activation in response to focal mechanical stress may be helpful in understanding mechanotransduction signaling.

## Materials and Methods

### cDNA construction

Hybrid cDNAs encoding fusions of bovine PKCα (Plasmid 10805; Invitrogen) and DG (pDG1-MN1, AM-V0073; Medical & Biological Laboratories Co., LTD, JAPAN) were inserted into the multiple cloning site of the pcDNATM3.1(+) (V790-20; Invitrogen) mammalian expression vector. We created the linker that connected PKCα to DG (Wagner et al., 2000). cDNAs encoding PKCα, DG, and the linker were amplified by polymerase chain reaction using the following primers: 5'-PKCα, 5'-ACCCAAGCTGGCTAGCACCATGGCTGAC-3'; 3'-PKCα, 5'-GTCGACTGCAGAATTCGCCCTTGTTTACTACCG-3'; 5'-DG, 5'-CCCGGGATCCACCGACCGGTCGCCACCATGGTGAGTGTGATT-3'; 3'-DG, 5'-GCCCTCTAGACTCGAGACTTGGCCTGCCTCGGCAG -3'; 5'-Linker, 5'-TAAGAATTCTGCAGTCGACGG-3'; and 3'-Linker, 5'-TAATATGCGGCCGCTATACC-3'. The NheI/EcoRI-digested PKCα fragment and the AgeI/XhoI-digested DG fragment were ligated into the NheI/XhoI-digested vector, generating the PKCα–DG fusion construct. All constructs were confirmed by DNA sequencing (Bio Matrix Research, Inc., Kashiwa, Chiba, Japan).

### Cell culture and transfection

BAECs (Toyobo Co., Ltd., Osaka, Japan) were cultured in Dulbecco’s Modified Eagle’s Medium (DMEM, GIBCO™; Thermo Fisher Scientific, Waltham, MA, USA), containing 1% Antibiotic-Antimycotic (10000 U/ml penicillin, 10000 μg/ml streptomycin, and 25 μg/ml amphotericin B, GIBCO™; Thermo Fisher Scientific) and 10% (v/v) fetal bovine serum (Biological Industries, Kibbutz Beit-Haemek, Israel), in humidified incubators containing 5% CO2 at 37 °C.

Prior to transfection, BAECs were seeded in a 35-mm glass-bottomed dish and cultured to 70% confluence. BAECs adherent to the glass-bottomed dishes were washed twice with opti-MEM (GIBCO™; Thermo Fisher Scientific). We changed DMEM for 1 ml of opti-MEM, and the BAECs were cultured for 14–16 h. Two microliters of DNA (1 μg/μl) in 100 μl of opti-MEM were mixed with 4 μl of HilyMax (Dojindo Laboratories, Kumamoto, Japan). This solution, including the construct DNA, was then added to the BAECs adherent to the glass-bottomed dish, together with 1 ml of opti-MEM, and PKCα–DG was transfected into the BAECs. We used BAECs (Passages 6–10) grown to confluence for experiments 2 days after transfection.

### Immunoblotting

After the protein concentration was measured using the Lowry method, equivalent amounts of protein were prepared and separated on a 7.5% polyacrylamide-sodium dodecyl sulfate gel and transferred to an Immun-Blot® polyvinylidene difluoride membrane (Bio-Rad Laboratories, Inc., Hercules, CA, USA). The membranes were washed three times with Tris-buffered saline (TBS;0.02 M Tris and 0.5 M NaCl; Bio-Rad Laboratories, Inc.) containing 0.1% (v/v) Tween-20 (Bio-Rad Laboratories, Inc.). The membranes were successively incubated, first with blocking buffer containing TBS, 0.1% (v/v) Tween-20, and 3% (w/v) Albumin (Nacalai Tesque, Inc., Kyoto, Japan) for 1 h at room temperature. The next incubation was conducted with the primary antibodies anti-PKCα Clone M4 (EMD Millipore Corp., Billerica, MA, USA) and anti-DG (MBL Laboratories Co, JAPAN) at a dilution of 1:5000 for 1 h at room temperature. After washing twice with TBS containing 0.1% (v/v) Tween-20, a third incubation was performed with the secondary antibody anti-mouse Immunoglobulin G (IgG) biotin (eBioscience, Inc., San Diego, CA, USA) at a dilution of 1:5000 for 1 h at room temperature. Thereafter, the membranes were probed with Pierce® High-Sensitivity Streptavidin-HRP (Thermo Fisher Scientific) at a dilution of 1:5000 for 1 h at room temperature. Finally, the membranes were incubated with Luminata Forte Western HRP (Merck KGaA) for 1 min at room temperature.

### Characterization of the PKCα–DG in BAECs

For immunocytochemistry, transfected BAECs were fixed with 4% paraformaldehyde for 10 min. After washing twice with phosphate-buffered saline (PBS), BAECs were permeabilized with PBS containing 0.1% (v/v) Triton-X for 5 min. After washing three times with PBS, the preparation was incubated for 1 h at room temperature with mouse anti-PKCα Clone M4 (Clontech Laboratories, Inc.) at a dilution of 1:200. The preparation was washed three times with PBS, and then exposed to the secondary antibody (Alexa Fluor 568-conjugated anti-mouse IgG, 1:200; Life Technologies, Carlsbad, CA, USA) for 1 h. Then, the preparation was washed three times with PBS, and observed in 4-(2-hydroxyethyl)-1-piperazineethanesulfonic acid (HEPES)-buffered saline (HBS; 140 mM NaCl, 5 mM KCl, 1 mM MgCl2, 1.8 mM CaCl2, 10 mM D-glucose, and 15 mM HEPES). Fluorescent images were captured by confocal microscopy (Digital Eclipse C1; Nikon Corp., Tokyo, Japan) using a 60X oil-immersion lens (S-Fluor, NA = 1.4; Nikon Corp.). Confocal images of DG were obtained by excitation with a sapphire laser at 488 nm and using a reflection short-pass 480-nm filter for GFP. Confocal images of Alexa Fluor 568 were obtained by excitation with a helium neon laser at 543 nm and using a reflection 545-nm filter. Emission wavelengths were detected using a 515 ± 15 nm filter for GFP and a 605 ± 30 nm filter for Alexa Fluor 568.

For pharmacological study, the translocation of PKCα–DG to the cell membrane was observed in the addition of 200 μM ATP and 500 μM PMA (Sigma-Aldrich Corp., St. Louis, MO, USA) using confocal microscopy (Digital Eclipse C1) with a 60X oil-immersion lens (S-Fluor, NA= 1.4). Confocal images of PKCα–DG were obtained by excitation with a sapphire laser at 488 nm and using a reflection short-pass 480 nm filter. The emission wavelength was detected using a 515± 15 nm filter.

### Mechanical stimulation

The mechanical stimulation of BAECs was performed on a single cell using a fine glass micropipette. The tip of the pipette was pulled so as to be less than 3 μm in diameter (P-97; Sutter Instrument Company, Novato, CA, USA). To prevent rupture of the plasma membrane, the tip of the micropipette was heat-polished. The micropipette was fixed in a micromanipulator (MMO-203; Narishige Group, Tokyo, Japan). The tip of the micropipette was softly attached to the plasma membrane surface. Micropipette indention depth was approximately 1 μm–3 μm, a depth which was estimated using the scale on the manual micromanipulator.

### Simultaneous measurement of changes in intracellular Ca^2+^ concentration and PKCα–DG translocation

BAECs were cultured in a 35-mm glass-bottomed dish to 100% confluence for 48 h after transfection. In this study, in addition to PKCα kinetics, we examined the intracellular Ca^2+^ wave in response to the microscopic mechanical stress by observing the decrease in the fluorescent intensity of the Ca^2+^ indicator Fura-2–acetoxymethyl ester (Fura-2–AM; Invitrogen, Carlsbad, CA, USA). Transfected BAECs were washed twice with HBS and then loaded with 5 μM Fura-2–AM in HBS for 30 min at room temperature. After washing twice with HBS, the transfected and Fura-2–AM-loaded cells were placed on the stage of an inverted microscope (Eclipse TE2000-S; Nikon Corp.) fitted with a 40X oil-immersion lens (S-Fluor, NA = 1.30; Nikon Corp.). In the light path, a 500 nm dichroic mirror and a 535 ± 45 nm emission filter were used. The excitation light was emitted by a xenon lamp (C7773; Hamamatsu Photonics K.K., Hamamatsu, Shizuoka Japan), and the wavelength was filtered using the AquaCosmos system (version 2.61; Hamamatsu Photonics K.K.).The Fura-2–AM was excited with 390 nm, and the PKCα–DG was excited with 490 nm. To monitor the fluorescence of both intracellular Ca^2+^ and PKCα–DG simultaneously, the excitation wavelength shifted at 148 ms intervals. The fluorescent emission from the sample was amplified with an image intensifier (C8600-03; Hamamatsu Photonics K.K.), detected by a charge-coupled device camera (C6790; Hamamatsu Photonics K.K.), and recorded on a personal computer. The microscope stage was maintained at 37 °C by enclosure in thermal insulation material.

In the inhibition experiments, the following inhibitors were used: 20 nM Bryostatin (Sigma-Aldrich Corp., St. Louis, MO, USA) for the DAG-dependent C1 domain for PKCα; and 10 μM Go6976 (Sigma-Aldrich Corp., St. Louis, MO, USA) for inhibiting the Ca^2+^-dependent C2 domain of PKCα. Additionally, to investigate the effect of Ca^2+^ on PKCα kinetics, 200 nM Thapsigargin (Sigma-Aldrich Corp., St. Louis, MO, USA) for inhibition of Ca^2+^ release from the ER, and 20 μM Gadolinium Chloride Hexahydrate (Gd^3+^; Wako pure chemical industries, Ltd, JAPAN) for inhibiting the MS channel were loaded onto the BAECs after Fura-2-AM loading. Gd^3+^ and other inhibitors were dissolved in HBS and DMSO, respectively. These inhibitors were loaded for 30 min for Gd^3+^ and Go6976, for 1 hour for Thapsigargin, and for 6 hours for Bryostatin, and then washed with HBS. Furthermore, for the removal of extracellular Ca^2+^ in HBS, HBS was exchanged with HBS containing EGTA (140 mM NaCl, 5 mM KCl, 1 mM MgCl2, 10 mM D-glucose, 15 mM HEPES, and 200μM EGTA).

PKCα–DG and intracellular Ca^2+^ fluorescent intensity were evaluated according to the intensity ratio in the region of interest (ROI) with a diameter of 2 μm. The fluorescent intensity ratio was defined as the fluorescent intensity at any time divided by the time-averaged intensity before stimulation during about 3 s (20 images). In this study, we measured the intensity of PKC-DG and Ca^2+^ at the stimulated point, at the cell membrane nearby, and in the cytoplasm, respectively. Regarding the cell membrane nearby, the increase in PKC-DG intensity was not homogenous throughout the membrane. Therefore, the average image before stimulation was subtracted from all the images, and the ROI was defined as the specific spot with the increasing intensity extending to a maximum depth of 4 μm from the edge of the cell. Regarding the cytoplasm, the ROI was defined as the area between the stimulation spot and the specific membrane spot.

### Statistical analysis

Statistical analyzes were performed using analysis of variance. Multiple comparisons were performed using Dunnett’s test. Differences were considered significant at a p value < 0.05. The number of glass-bottomed culture dishes and cells were denoted by N and n, respectively.

## Conflict of Interest Statement

No conflicts of interest, financial or otherwise, are declared by the authors.

### Acknowledgments

This research was partially supported by JSPS KAKENHI Grant Number JP16H02529.

## References

Bazzi, M. D. and Nelsestuen, G. L. (1989). Differences in the effects of phorbol esters and diacylglycerols on protein kinase C. Biochemistry 28, 9317–9323.

Bokhari, S. M., Zhou, L., Karasek, M. A., Paturi, S. G. and Chaudhuri, V. (2006). Regulation of skin microvasculature angiogenesis, cell migration, and permeability by a specific inhibitor of PKCalpha. J. Invest. Dermatol. 126, 460–467.

Erickson, G. R., Alexopoulos, L. G. and Guilak, F. (2001). Hyper-osmotic stress induces volume change and calcium transients in chondrocytes by transmembrane, phospholipid, and G-protein pathways. J Biomech 34, 1527–1535.

Grygorczyk, R., Furuya, K. and Sokabe, M. (2013). Imaging and characterization of stretch-induced ATP release from alveolar A549 cells. J. Physiol. (Lond.) 591, 1195–1215.

Gudi, S., Nolan, J. P. and Frangos, J. A. (1998). Modulation of GTPase activity of G proteins by fluid shear stress and phospholipid composition. Proc. Natl. Acad. Sci. U.S.A. 95, 2515– 2519.

Hayakawa, K., Tatsumi, H. and Sokabe, M. (2008). Actin stress fibers transmit and focus force to activate mechanosensitive channels. J. Cell. Sci. 121, 496–503.

Hu, Y., Böck, G., Wick, G. and Xu, Q. (1998). Activation of PDGF receptor alpha in vascular smooth muscle cells by mechanical stress. FASEB J. 12, 1135–1142.

Ishida, T., Takahashi, M., Corson, M. A. and Berk, B. C. (1997). Fluid Shear Stress-Mediated Signal Transduction: How Do Endothelial Cells Transduce Mechanical Force into Biological Responses? Annals of the New York Academy of Sciences 811, 12–24.

Kazanietz, M. G., Krausz, K. W. and Blumberg, P. M. (1992). Differential irreversible insertion of protein kinase C into phospholipid vesicles by phorbol esters and related activators. J. Biol. Chem. 267, 20878–20886.

Konopatskaya, O. and Poole, A. W. (2010). Protein kinase Calpha: disease regulator and therapeutic target. Trends Pharmacol. Sci. 31, 8–14.

Lenz, J. C., Reusch, H. P., Albrecht, N., Schultz, G. and Schaefer, M. (2002). Ca2+-controlled competitive diacylglycerol binding of protein kinase C isoenzymes in living cells. J. Cell Biol. 159, 291–302.

Li, C. and Xu, Q. (2000). Mechanical stress-initiated signal transductions in vascular smooth muscle cells. Cellular Signalling 12, 435–445.

Li, C. and Xu, Q. (2007). Mechanical stress-initiated signal transduction in vascular smooth muscle cells in vitro and in vivo. Cellular Signalling 19, 881–891.

Li, C., Wernig, F., Leitges, M., Hu, Y. and Xu, Q. (2003). Mechanical stress-activated PKCdelta regulates smooth muscle cell migration. FASEB J. 17, 2106–2108.

Luo, J. H. and Weinstein, I. B. (1993). Calcium-dependent activation of protein kinase C. The role of the C2 domain in divalent cation selectivity. J. Biol. Chem. 268, 23580–23584.

Ma, Y.-H., Ling, S. and Ives, H. E. (1999). Mechanical Strain Increases PDGF-B and PDGF β Receptor Expression in Vascular Smooth Muscle Cells. Biochemical and Biophysical Research Communications 265, 606–610.

Maasch, C., Wagner, S., Lindschau, C., Alexander, G., Buchner, K., Gollasch, M., Luft, F. C. and Haller, H. (2000). Protein kinase calpha targeting is regulated by temporal and spatial changes in intracellular free calcium concentration [Ca(2+)](i). FASEB J. 14, 1653–1663.

Medkova, M. and Cho, W. (1999). Interplay of C1 and C2 Domains of Protein Kinase C-α in Its Membrane Binding and Activation. J. Biol. Chem. 274, 19852–19861.

Moriya, S., Kazlauskas, A., Akimoto, K., Hirai, S., Mizuno, K., Takenawa, T., Fukui, Y., Watanabe, Y., Ozaki, S. and Ohno, S. (1996). Platelet-derived growth factor activates protein kinase C epsilon through redundant and independent signaling pathways involving phospholipase C gamma or phosphatidylinositol 3-kinase. Proc Natl Acad Sci U S A 93, 151–155.

Nalefski, E. A. and Newton, A. C. (2001). Membrane Binding Kinetics of Protein Kinase C βII Mediated by the C2 Domain. Biochemistry 40, 13216–13229.

Naruse, K. and Sokabe, M. (1993). Involvement of stretch-activated ion channels in Ca2+ mobilization to mechanical stretch in endothelial cells. Am. J. Physiol. 264, C1037–1044.

Newton, A. C. (2001). Protein Kinase C:? Structural and Spatial Regulation by Phosphorylation, Cofactors, and Macromolecular Interactions. Chem. Rev. 101, 2353–2364.

Oancea, E. and Meyer, T. (1998). Protein kinase C as a molecular machine for decoding calcium and diacylglycerol signals. Cell 95, 307–318.

Schaefer, M., Albrecht, N., Hofmann, T., Gudermann, T. and Schultz, G. (2001). Diffusion-limited translocation mechanism of protein kinase C isotypes. FASEB J. 15, 1634–1636.

Sinnecker, D. and Schaefer, M. (2004). Real-time analysis of phospholipase C activity during different patterns of receptor-induced Ca2+ responses in HEK293 cells. Cell Calcium 35, 29–38.

Takada, H., Furuya, K. and Sokabe, M. (2014). Mechanosensitive ATP release from hemichannels and Ca^2+^ influx through TRPC6 accelerate wound closure in keratinocytes. J. Cell. Sci. 127, 4159–4171.

Takahara, N., Ito, S., Furuya, K., Naruse, K., Aso, H., Kondo, M., Sokabe, M. and Hasegawa, Y. (2014). Real-time imaging of ATP release induced by mechanical stretch in human airway smooth muscle cells. Am. J. Respir. Cell Mol. Biol. 51, 772–782.

Tanabe, Y., Saito, M., Ueno, A., Nakamura, M., Takeishi, K. and Nakayama, K. (2000). Mechanical stretch augments PDGF receptor β expression and protein tyrosine phosphorylation in pulmonary artery tissue and smooth muscle cells. Mol Cell Biochem 215, 103–113.

Tsukamoto, A., Hayashida, Y., Furukawa, K. S. and Ushida, T. (2010). Spatio-temporal PLC activation in parallel with intracellular Ca2+ wave propagation in mechanically stimulated single MDCK cells. Cell Calcium 47, 253–263.

Wagner, S., Harteneck, C., Hucho, F. and Buchner, K. (2000). Analysis of the Subcellular Distribution of Protein Kinase Cα Using PKC-GFP Fusion Proteins. Experimental Cell Research 258, 204–214.

Wang, A., Nomura, M., Patan, S. and Ware, J. A. (2002). Inhibition of protein kinase Calpha prevents endothelial cell migration and vascular tube formation in vitro and myocardial neovascularization in vivo. Circ. Res. 90, 609–616.

Yellowley, C. E., Jacobs, C. R. and Donahue, H. J. (1999). Mechanisms contributing to fluid-flow-induced Ca2+ mobilization in articular chondrocytes. J. Cell. Physiol. 180, 402–408.

